# DHHC3-dependent S-Acylation of CRY1 regulates its subcellular localization and repressor function in the circadian clock

**DOI:** 10.64898/2026.05.20.726662

**Authors:** Ji Ye Lim, Shayahati Bieerkehazhi, Chorong Han, Sun Young Kim, Matthew L. Baker, Tingting Mills, Kuang-Lei Tsai, Hyun Kyoung Lee, Sung Yun Jung, Zheng Chen, Askar M. Akimzhanov, Seung-Hee Yoo

## Abstract

The circadian clock is essential for maintaining cellular homeostasis and physiological fitness. At the molecular level, core clock proteins function via transcriptional-translational feedback loops in the cellular oscillator, and are highly regulated by post-translational modifications. Our unbiased screening of core clock proteins revealed that Cryptochrome 1 (CRY1), the central transcriptional repressor in the circadian clock, undergoes a novel post-translational modification known as S-acylation. We show that this reversible lipidation of CRY1 is required for its nuclear import and interaction with key clock components. Further, we mapped four cysteine residues as CRY1 S-acylation sites and identified DHHC3 as the primary protein acyltransferase for CRY1. Importantly, loss of CRY1 S-acylation, either via cysteine mutagenesis or genetic deletion of DHHC3, impaired CRY1 repressor function and consequently cellular circadian rhythms, suggesting that dynamic S-acylation couples cytoplasmic regulation of CRY1 and its transcriptional repressor function in the nucleus. Together, our findings identify S-acylation as a previously unknown post-translational modification of CRY1 critical for circadian clock function and establish DHHC3 as a pivotal circadian regulatory enzyme. Targeting CRY1 S-acylation or its regulatory enzymes may constitute an innovative therapeutic approach against clock-associated diseases.

## Introduction

The circadian clock is our intrinsic biological timer composed of cellular oscillators in almost all cells in our body and regulates tissue-specific oscillatory gene expression to control metabolism, physiology, and behavior (Bass and Lazar, 2016; Takahashi, 2017). Disruptions of the circadian clock are associated with a broad range of pathologies, including metabolic diseases, obesity, immune system disorders and cancer (Bass and Takahashi, 2010; Cederroth et al., 2019; Masri and Sassone-Corsi, 2018). At the molecular level, the oscillator is driven by transcriptional-translational feedback loops containing positive (CLOCK, NPAS2, BMAL1, RORs) and negative (PER1/2, CRY1/2, REV-ERBs) core clock components (Green et al., 2008; Mure et al., 2018; Takahashi, 2017; Zhang et al., 2014). Importantly, post-translational modifications (PTMs), such as phosphorylation and ubiquitination, of clock proteins play crucial roles in establishing robust circadian rhythmicity and enabling timely responses to environmental cues (Gallego and Virshup, 2007; Hirano et al., 2016; Millius et al., 2019; Okamoto-Uchida et al., 2019).

CRY1 and CRY2 serve as essential transcriptional repressors in the oscillator to inhibit CLOCK– BMAL1 activity and thereby dictate the amplitude and precision of circadian rhythms (Kume et al., 1999; van der Horst et al., 1999; Vitaterna et al., 1999). The stability, abundance, and subcellular distribution of CRY proteins are tightly controlled. Multiple post-translational modifications, including phosphorylation and ubiquitination, have been shown to modulate the function of CRY proteins and circadian gene expression (Hirano et al., 2013; Kurabayashi et al., 2010; Lamia et al., 2009; Yoo et al., 2013). For example, we and others have shown that the CRY proteins are regulated by two antagonistically acting circadian E3 ubiquitin ligases, FBXL3 and FBXL21 (Hirano et al., 2013; Yoo et al., 2013). We found that loss of FBXL3 function results in a lengthened circadian period, whereas a hypomorph mutation of *Fbxl21* shortens the period (Siepka et al., 2007; Yoo et al., 2013). Mechanistically, FBXL21 slowly degrades CRY proteins in the cytoplasm but antagonizes the stronger E3 ligase activity of FBXL3 to stabilize CRY in the nucleus (Yoo et al., 2013). These findings highlight the importance of spatial compartmentalization in regulating CRY stability and protein-protein interactions, factors that critically determine the circadian period in mammals. Yet, despite extensive studies, the molecular mechanism controlling CRY intracellular localization and dynamic interactions with binding partners is not fully understood.

Protein S-acylation (also termed S-palmitoylation) is a post-translational modification of cysteine residues with long-chain fatty acids via a labile thioester bond (Chamberlain and Shipston, 2015). Unlike other types of protein lipidation such as N-myristoylation or prenylation, S-acylation is fully reversible, and the modified proteins can undergo multiple cycles of acylation/de-acylation (Chamberlain and Shipston, 2015; Jiang et al., 2018; Zaballa and van der Goot, 2018). This unique property of S-acylation allows for rapid change in protein hydrophobicity to regulate multiple aspects of protein function, including localization to membranes, intracellular trafficking, and protein-protein interactions (Chen et al., 2021; West et al., 2022). Advances in proteomic methods have revealed that up to 20% of the human proteome may be subject to S-acylation, highlighting its widespread physiological relevance (Anwar and van der Goot, 2023; Chen et al., 2018; F et al., 2024). Due to its high lability and prominent impact on intracellular localization, S-acylation is uniquely positioned as a dynamic switch regulating protein distribution within the cell. However, the role of protein S-acylation in circadian biology remains largely unexplored.

Here, we identify Cryptochrome 1 (CRY1) as an S-acylated protein and reveal a previously unrecognized layer of post-translational control within the circadian oscillator. To explore a possible role of reversible protein lipidation in the regulation of the circadian clock, we used the Acyl-Resin Assisted Capture (Acyl-RAC) assay to screen the core clock components and identified CRY1 as a novel S-acylated protein. Using site-specific Acyl-RAC proteomics, we mapped S-acylation sites on CRY1 and generated an acylation-deficient mutant to examine the functional consequences of this modification. We found that S-acylation of CRY1 was required for its nuclear translocation and interaction with PER2 and BMAL1, and that loss of this modification impaired CRY1’s ability to inhibit CLOCK:BMAL1-driven transcription. S-acylation of CRY1 was mainly catalyzed by the acyltransferase DHHC3, and that this modification was required for efficient nuclear import, assembly into repressor complexes, and robust circadian oscillation. Loss of S-acylation, either by mutating the modified cysteine residues or by deleting DHHC3, disrupted CRY1 repressor activity and attenuates cellular circadian rhythms. These findings establish protein S-acylation as a critical regulator of CRY1 and identify DHHC3 as a previously unrecognized enzymatic regulator of the circadian system.

## Results

### CRY1 is a novel S-acylated protein

To examine whether S-acylation contributes to the regulation of the mammalian circadian clock, we first assessed S-acylation of core clock proteins in murine tissues using the Acyl-Resin Assisted Capture (Acyl-RAC) Assay, previously optimized by our group (Tewari et al., 2020). Briefly, following cell or tissue lysis, free cysteine thiols (-SH) were irreversibly blocked by N-ethylmaleimide (NEM) to prevent unspecific detection (Fig. 1A). Thioester bonds between cysteines and acyl groups were then specifically cleaved by neutral hydroxylamine (NH_2_OH) (Fig. 1A). The newly formed free thiol groups were directly captured by S3 Thiol Sepharose resin and analyzed by immunoblotting to determine S-acylation of the target protein (Fig. 1A). We expressed the major core clock proteins (CLOCK, BMAL1, PER1, PER2, CRY1, CRY2, REV-ERBα, and RORα) in 293T cells and performed Acyl-RAC assays (Fig. 1B). Among these, only CRY1 exhibited detectable S-acylation, whereas no S-acylation was observed for any of the other core clock proteins (Fig. 1B). Next, we examined whether CRY1 is S-acylated *in vivo*. The S-acylation of CRY1 was detected in the liver, spleen (splenocytes), and cerebellum of wild-type (WT) mice (Figs. 1C-F and S1A). Notably, we observed robust circadian oscillations of CRY1 S-acylation, with peak levels occurring between Zeitgeber Time (ZT) 20 and ZT24 (ZT0) in both liver (Figs. 1C-D) and splenocytes (Figs. 1E-F), determined by JTK_Cycle analysis (Hughes et al., 2010). Consistent with our *in vitro* result (Fig. 1B), no S-acylation of CRY2 was detectable in the liver (Fig. S1B). These results indicate that CRY1 S-acylation is under circadian control and occurs in various tissues, suggesting a potential broad role of protein S-acylation in regulating time-of-day-dependent CRY1 function.

**Fig. 1.**
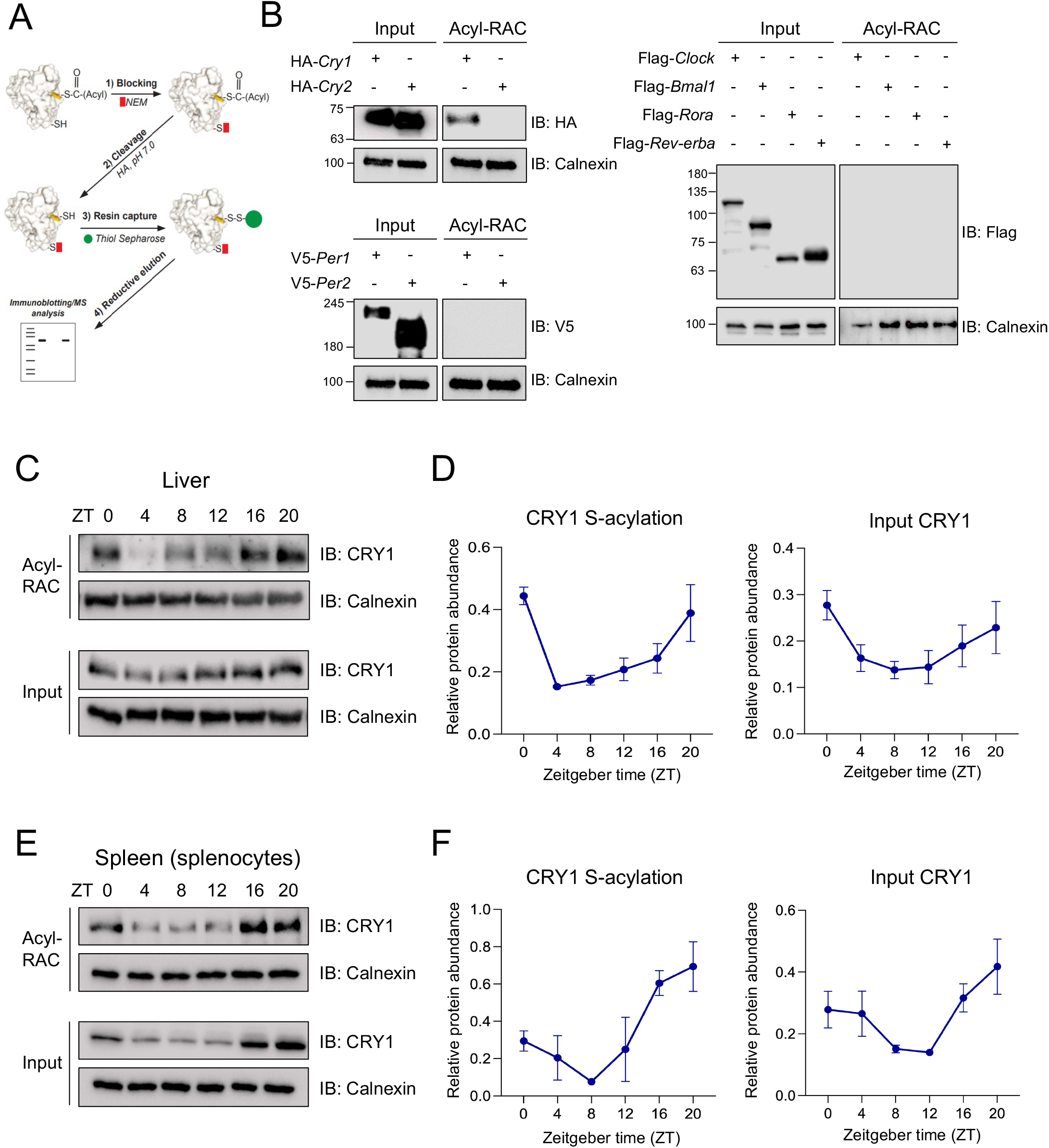
CRY1 is identified as an S-acylated core clock protein. (**A**) Schematic workflow of the Acyl-RAC assay. (**B**) Screening of core clock proteins (CRY1, CRY2, PER1, PER2, CLOCK, BMAL1, RORα, and REV-ERBα) for S-acylation using the Acyl-RAC assay. Calnexin, a known S-acylated protein, was used as a loading control. (**C**) Immunoblot analysis of CRY1 S-acylation in the liver tissues of WT C57BL/6J mice collected at the indicated Zeitgeber times (ZT), as determined by the Acyl-RAC assay. (**D**) Quantification of S-acylated CRY1 (left panel) and total input CRY1 (right panel) in liver tissues across circadian time points. Data are presented as mean ± SEM (n = 3 per time point). JTK_Cycle analysis showed that S-acylated CRY1 and input CRY1 are rhythmic in the liver tissues of WT mice (JTK_Cycle, adjusted p-value: \*\**p* < 0.01 for S-acylated CRY1 and \**p* < 0.05 for input CRY1). (**E**) Immunoblot analysis of CRY1 in splenocytes from WT mice collected at circadian time points. (**F**) Quantification of S-acylated CRY1 (left panel) and total input CRY1 (right panel) in splenocytes across circadian time points. Data are presented as mean ± SEM (n = 3 per time point). JTK_Cycle analysis showed that S-acylated CRY1 and input CRY1 are rhythmic in splenocytes from WT mice (JTK_Cycle, adjusted p-value: \*\**p* < 0.01 for S-acylated CRY1 and \**p* < 0.05 for input CRY1).

### S-acylation of CRY1 is required for its nuclear translocation and cellular circadian rhythms by regulating interaction with other core clock proteins

To identify the modified cysteine residue(s), we integrated Acyl-RAC with mass spectrometry (MS) to analyze Cys-containing peptides selectively retained on the thiol-reactive resin (Fig. 2A). Briefly, we transfected HEK293T cells with Strep-tagged-CRY1. Following cell lysis and neutral hydroxylamine treatment, S-acylated proteins were captured with thiol-reactive resin (Fig. 2A). Unlike the standard Acyl-RAC workflow, immobilized proteins were digested directly on the beads using trypsin, and the unbound peptides were removed by extensive washing. The retained peptides were then eluted by reducing disulfide bonds using DTT and subjected to LC-MS/MS analysis (Fig. 2A). MS analysis of captured peptides identified S-acylated proteome, including CRY1. Peptide mapping of CRY1 revealed four cysteines (Cys24, Cys58, Cys259, and Cys315) within the photolyase-homologous region of CRY1 as sites of S-acylation (Fig. 2A). Of note, these sites are not disulfide-bond–forming cysteines in CRY1, reinforcing that the detected modification represents true S-acylation rather than thiol oxidation. To validate our findings, we next generated a CRY1 mutant (4Cm-CRY1) in which identified cysteines were substituted with alanine (C24A, C58A, C259A, C315A) and performed Acyl-RAC analysis. Acyl-RAC analysis demonstrated more than 90% reduction in S-acylation levels in 4Cm-CRY1 compared to wild-type CRY1 (WT-CRY1) (Fig. 2B), confirming that these residues represent the primary S-acylation sites in CRY1. Next, to examine the structural impact and energetic consequences of the CRY1 S-acylation mutations in 4Cm-CRY1, we performed structural and computational analyses using AlphaFold3 modeling and Rosetta-based mutational analyses. Mapping these residues to the CRY1 structure (PDBID: 4K0R)(Czarna et al., 2013) indicated that the mutations in 4Cm-CRY1 do not disrupt global folding or stability (Fig. S2B).

**Fig. 2.**
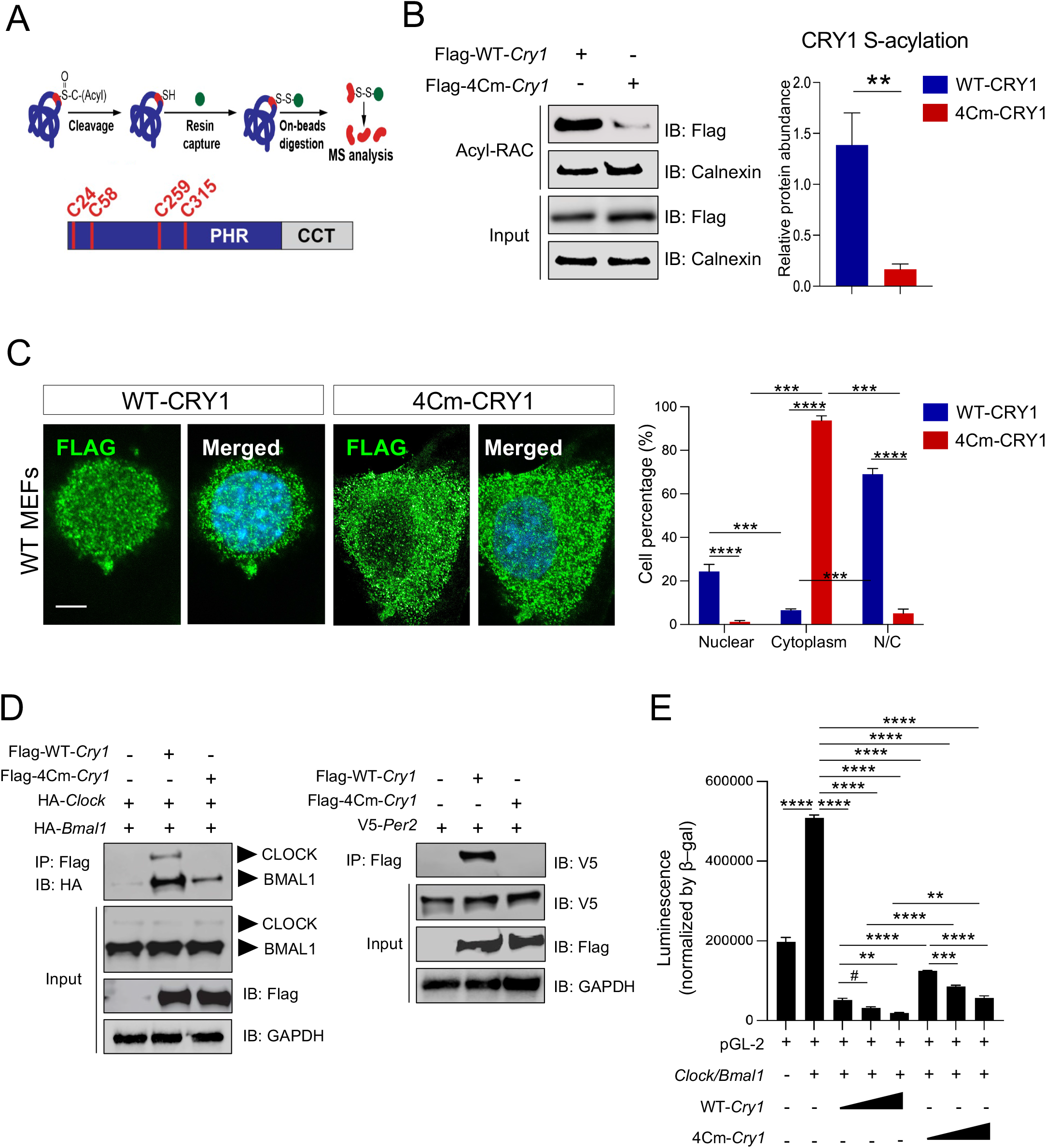
Identification of CRY1 S-acylation sites and the functional impact on core clock protein interaction, nuclear translocation and cellular circadian rhythms. (**A**) Schematic workflow of LC/MS analysis used to identify S-acylation sites on CRY1. (**B**) Immunoblot analysis of Acyl-RAC and input samples of flag-tagged WT-CRY1 and 4Cm-CRY1. Right panel: quantification of CRY1 S-acylation. Data are presented as mean ± SEM (n = 5). \*\**p* < 0.01; Student’s *t*-test indicates a significant difference between WT-CRY1 and 4Cm-CRY1. (**C**) STED microscopy analysis of FLAG expression in WT MEFs transiently transfected with Flag-tagged WT-*Cry1* or 4Cm-*Cry1*. Right panels: Quantification of the percentage of cells showing nuclear, cytoplasmic, or mixed nuclear/cytoplasmic localization. Data are presented as mean ± SEM (n = 3). \*\**p* < 0.01, \*\*\**p* < 0.001, and \*\*\*\**p* < 0.0001; two-way ANOVA indicates statistical differences between groups. Scale bar: 5 μm. (**D**). The interaction between BMAL1/CLOCK, PER2, and WT-CRY1 or 4Cm-CRY1. HEK293T cells were co-transfected with the indicated plasmids, and protein complexes were pulled down using anti-Flag beads. (**E**) Effects of WT-CRY1 or 4Cm-CRY1 on CLOCK/BMAL1-mediated transcriptional activation. pGL-2 Per2 promoter and indicated plasmids were transfected into 293T cells and luciferase activity was measured. Data are presented as mean ± SEM (n = 3–4). \*\**p* < 0.01, \*\*\*\*\**p* < 0.001, and \*\*\*\**p* < 0.0001; One-way ANOVA indicates statistical differences between groups. ^#^*p* < 0.05; Student’s *t*-test indicates a significant difference between groups.

Translocation into the nucleus is vital for CRY1 function as the primary transcriptional repressor in the core oscillator. To assess the impact of S-acylation on CRY1 subcellular distribution, we used stimulated emission depletion (STED) microscopy to visualize Flag-tagged WT-CRY1 and the acylation-deficient 4Cm-CRY1 mutant transiently expressed in WT MEFs. Consistent with previous reports (Chaves et al., 2006; Miyazaki et al., 2001; Tamanini et al., 2005; van der Schalie et al., 2007), WT-CRY1 predominantly localized to the nucleus in WT MEFs (Fig. 2C). In contrast, 4Cm-CRY1 exhibited significantly reduced nuclear accumulation and increased cytoplasmic localization (Fig. 2C), suggesting an important role of S-acylation for efficient nuclear import of CRY1.

Direct protein-protein interactions with core clock components are essential for CRY1 repressor function in the circadian clock (Parico et al., 2020; Takahashi, 2017). To investigate whether S-acylation affects the ability of CRY1 to bind its known partners, we performed a series of co-immunoprecipitation (Co-IP) assays using Flag-tagged WT-CRY1 and 4Cm-CRY1 expressed in HEK293T cells. First, we co-expressed WT or acylation-deficient 4Cm-CRY1 with BMAL1 and CLOCK proteins. While WT-CRY1 strongly co-immunoprecipitated with BMAL1 in the presence of CLOCK, 4Cm-CRY1 exhibited significantly reduced binding (Fig. 2D). Next, we examined whether S-acylation affects CRY1 interaction with its heterodimerization partner, PER2. Unlike BMAL1 which binds CRY1 within the nucleus, CRY1 association with PER2 starts in the cytoplasm to form the heterodimer, where the interaction stabilizes CRY1 and promotes translocation of the CRY1:PER2 complex into the nucleus (Tamanini et al., 2005; Yagita et al., 2002). Interestingly, we found that the 4Cm-CRY1 mutant binding to PER2 was abolished compared to WT-CRY1 (Fig. 2D). The diminished nuclear translocation and impaired core clock proteins binding of the acylation-deficient mutant suggest that S-acylation of CRY1 plays a role in regulating its transcriptional repressor activity and circadian rhythms. To assess the effect of S-acylation on CRY1’s transcription repressor function on E-box, we performed transcription reporter assays using a pGL-2 luciferase reporter driven by a Per2 E’ box-containing promoter (Yoo et al., 2005). In HEK293T cells, WT-CRY1 suppressed CLOCK:BMAL1-driven luciferase activity in a concentration-dependent manner (Fig. 2E). In contrast, 4Cm-CRY1 displayed significantly impaired repressor activity compared to WT-CRY1 (Figs. 2E and S2). Together, these results indicate that loss of S-acylation impairs the nuclear localization and weakens protein-protein interactions of CRY1, resulting in diminished transcriptional repressor activity.

### S-Acylation of CRY1 is required for FBXL3/FBXL21-mediated ubiquitination and degradation

S-acylation has been shown to regulate ubiquitin-mediated protein turnover by controlling protein localization and stability, and disruption of S-acylation often leads to enhanced ubiquitination and degradation (Chamberlain and Shipston, 2015). As mentioned above, CRY1 stability is controlled by the E3 ubiquitin ligases FBXL3 and FBXL21, in a cell compartment-specific manner (Yoo et al., 2013). FBXL3 primarily promotes CRY1 degradation within the nucleus, while FBXL21 functions predominantly in the cytoplasm (Yoo et al., 2013). To investigate whether inhibition of CRY1 S-acylation alters ubiquitination by FBXL3 and FBXL21, we first assessed the ability of FBXL3 and FBXL21 to interact with 4Cm-CRY1. Compared with the significantly reduced interaction with core clock transcription components, 4Cm-CRY1 mutant showed relatively strong binding with both FBXL3 and FBXL21, albeit to a lesser extent than WT-CRY1 (Figs. 3A and S3). To assess degradation by FBXL3 and FBXL21, we measured CRY1 half-life using cycloheximide (CHX) assays. Whereas ectopic expression of FBXL3 in 293T cells accelerated WT-CRY1 degradation, it had little effect on 4Cm-CRY1 turnover (Figs. 3B and S3). Interestingly, FBXL21 markedly accelerated 4Cm-CRY1 degradation compared to WT-CRY1 (Figs. 3B and S3). These results suggest that inhibition of CRY1 S-acylation increased cytoplasmic localization (Fig. 2C) and induced a functional switch from FBXL3-mediated nuclear degradation to FBXL21-mediated cytoplasmic degradation. Next, we performed *in vitro* ubiquitination assays using WT- and 4Cm-CRY1 in the presence of FBXL3 or FBXL21. Consistent with the degradation results (Fig. 3B), FBXL3 primarily ubiquitinated WT-CRY1, whereas FBXL21 preferentially ubiquitinated 4Cm-CRY1 (Fig. 3C). Together, these findings indicate that CRY1 S-acylation is a critical regulatory step governing compartment-specific ubiquitination and degradation by FBXL3 and FBXL21.

**Fig. 3.**
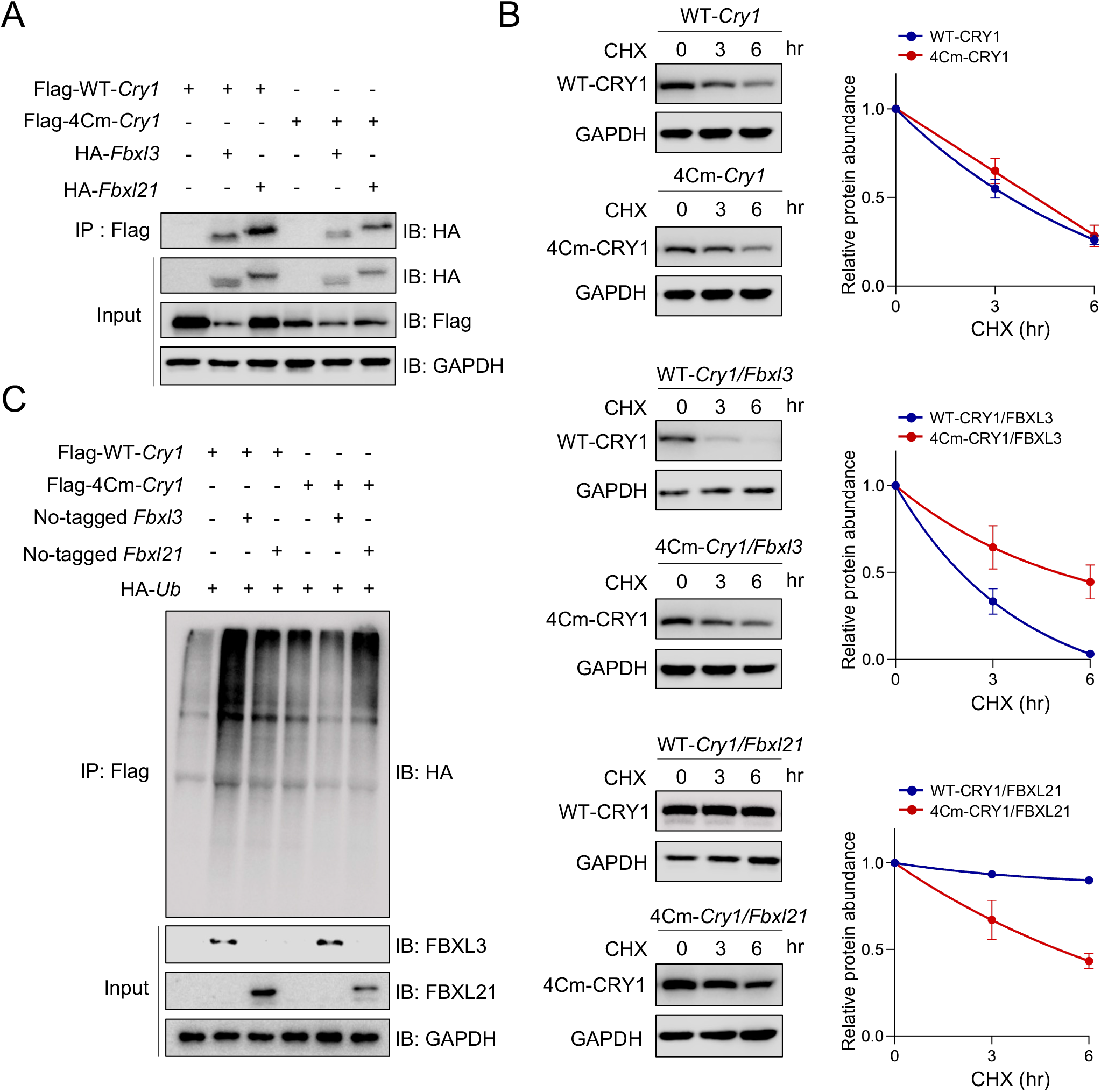
CRY1 S-acylation is required for FBXL3/FBXL21-mediated ubiquitination and degradation. (**A**) Interaction between CRY1 (WT-CRY1 or 4Cm-CRY1) and FBXL3 or FBXL21 was examined in HEK293T cells co-transfected with the indicated plasmids. Protein complexes were pulled down using anti-Flag beads. (**B**) Cycloheximide (CHX) chase assays were performed to assess the protein stability of WT-CRY1 and 4Cm-CRY1 in 293T cells. 293T cells were co-transfected with the indicated plasmids. Immunoblots showing WT-CRY1 and 4Cm-CRY1 protein stability. Half-lives were calculated using nonlinear one-phase decay analysis in GraphPad Prism. Half-lives: WT-Cry1, 3.25 h; 4Cm-Cry1, 3.77 h; WT-Cry1/Fbxl3, 1.72 h; 4Cm-Cry1/Fbxl3, 5.02 h; WT-Cry1/Fbxl21, >24.0 h; and 4Cm-Cry1/Fbxl21, 5.04 h. The decay constant (K) is significantly different between WT-Cry1/Fbxl3 and 4Cm-Cry1/Fbxl3 (\*\*\**p* < 0.001), as well as between WT-Cry1/Fbxl21 and 4Cm-Cry1/Fbxl21 (\*\*\*\**p* < 0.0001). Data are presented as mean ± SEM (n = 3). (**C**) Ubiquitination of 4Cm-CRY1 is not affected by FBXL3. Flag-tagged WT-Cry1 and 4Cm-Cry1 were co-transfected with untagged Fbxl3 or Fbxl21, and HA-tagged ubiquitin (HA-Ub), followed by ubiquitination assays.

### The protein acyltransferase DHHC3 mediates S-acylation of CRY1

In mammalian cells, S-acylation is catalyzed by the DHHC family of protein acyltransferases, which are named for their highly conserved Asp-His-His-Cys (DHHC) tetrapeptide motif within their catalytic core (Chamberlain and Shipston, 2015). The family comprises 23 members all of which are multi-pass transmembrane proteins, with the N- and C-termini present in the cytosol (Mitchell et al., 2006). To identify the enzyme responsible for S-acylation of CRY1, we conducted an unbiased Co-IP screen in which individual HA-tagged DHHC enzymes (encoded by mouse *Zdhhc* genes, generous gift from Dr. Masaki Fukata) (Fukata et al., 2006; Tsutsumi et al., 2009) were co-expressed with Flag-WT-CRY1 in HEK293T cells, and binding was assessed by immunoblotting. Among the members of the murine DHHC family, HA-tagged DHHC3, 17, 18, 19, and 22 were found to associate with Flag-tagged CRY1 (Fig. S4). Among these enzymes, HA-DHHC3 showed the strongest interaction with Flag-WT-CRY1 (Figs. 4A and S4), and, therefore, was selected as a leading candidate for further studies.

**Fig. 4.**
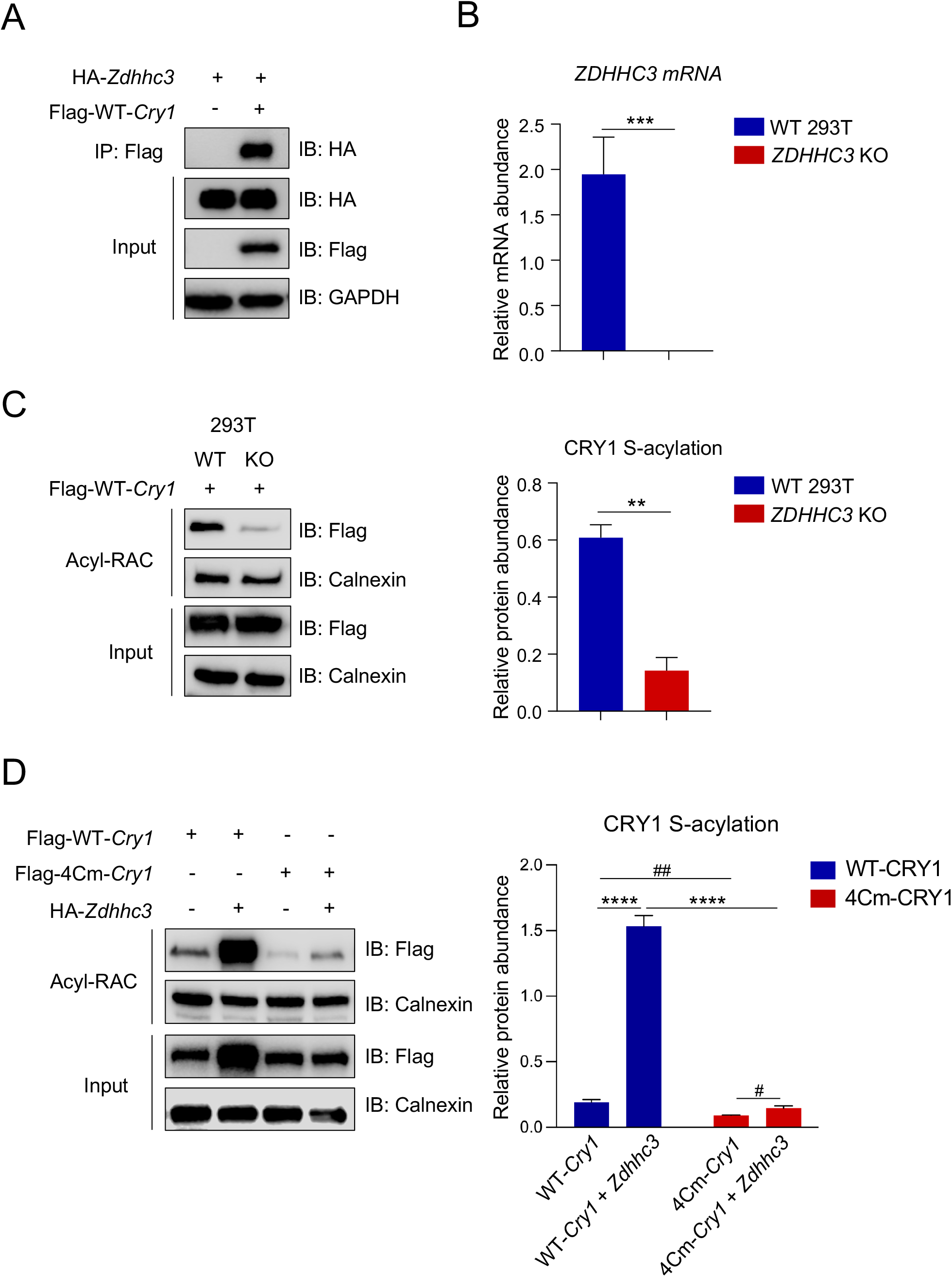
CRY1 S-acylation is mainly mediated by the protein acyltransferase DHHC3. (**A**) Interaction between DHHC3 and WT-CRY1. HEK293T cells were co-transfected with HA-tagged *Zdhhc3* and Flag-tagged WT-*Cry1*, and protein complexes were pulled down using anti-Flag beads. (**B**) Knockdown of *ZDHHC3* in *ZDHHC3* KO 293T cells. The mRNA levels of *ZDHHC3* were measured by qPCR in WT and *ZDHHC3* KO 293T cells. Data are presented as mean ± SEM (n = 3). \*\*\**p* < 0.001; Student’s *t*-test indicates a significant difference between WT and *ZDHHC3* KO 293T cells. (**C**) Reduced CRY1 S-acylation levels in *ZDHHC3* KO 293T cells. Left panel: Immunoblot analysis of Acyl-RAC and input samples from WT and *ZDHHC3* KO 293T cells expressing Flag-tagged WT-CRY1. Right panel: Quantification of WT-CRY1 S-acylation levels in WT and *ZDHHC3* KO 293T cells. Data are presented as mean ± SEM (n = 3). \*\**p* < 0.01; Student’s *t*-test indicates a significant difference between WT-CRY1 and 4Cm-CRY1. (**D**) Increased CRY1 S-acylation in the presence of DHHC3. Left panel: Representative immunoblots of Acyl-RAC and input samples from 293T cells expressing Flag-tagged WT-CRY1, 4Cm-CRY1, in the presence or absence of DHHC3 expression. Right panel: Quantification of CRY1 S-acylation levels. Data are presented as mean ± SEM (n = 3). \*\*\*\**p* < 0.0001; two-way ANOVA indicates statistical differences between groups. ^#^*p* < 0.05 and ^##^*p* < 0.01; Student’s *t*-test indicates a significant difference between groups.

To determine whether DHHC3 is required for CRY1 S-acylation, we generated a stable DHHC3-deficient 293T cell line using CRISPR–Cas9–mediated gene disruption. Loss of endogenous *ZDHHC3* expression in *ZDHHC3* KO 293T cells was confirmed by qPCR (Fig. 4B). Acyl-RAC assays revealed a marked reduction in S-acylation of Flag-WT-CRY1 in *ZDHHC3* KO compared to WT 293T cells (Fig. 4C), demonstrating that DHHC3 is necessary for S-acylation of CRY1. To investigate whether DHHC3 gain-of-function promotes S-acylation of CRY1, we performed Acyl-RAC assays in 293T cells expressing Flag-WT-CRY1 or Flag-4Cm-CRY1 in the presence or absence of DHHC3. Notably, DHHC3 expression significantly increased the S-acylation of WT-CRY1 (by 8.01-fold), compared to only a marginal effect on 4Cm-CRY1 (by 1.62-fold), indicating that these cysteine residues are the primary targets of DHHC3-mediated acylation. Furthermore, input WT-CRY1 protein levels were increased in the presence of DHHC3, suggesting that DHHC3-mediated S-acylation enhances WT-CRY1 stability. Together, these results suggest that DHHC3 is the primary acyltransferase for CRY1 with an important a role in CRY1 stability.

### DHHC3 regulates CRY1 localization and cellular circadian rhythms

To investigate whether DHHC3 deletion affects the subcellular localization of CRY1, we performed STED microscopy in *ZDHHC3* KO 293T cells expressing Flag-tagged WT-CRY1 or 4Cm-CRY1. Notably, DHHC3 deficiency significantly reduced nuclear localization of WT-CRY1, with a corresponding increase in cytoplasmic localization. (Fig. 5A). In contrast, 4Cm-CRY1 was predominantly cytoplasmic in both WT and *ZDHHC3* KO 293T cells (Fig. 5A).

**Fig. 5.**
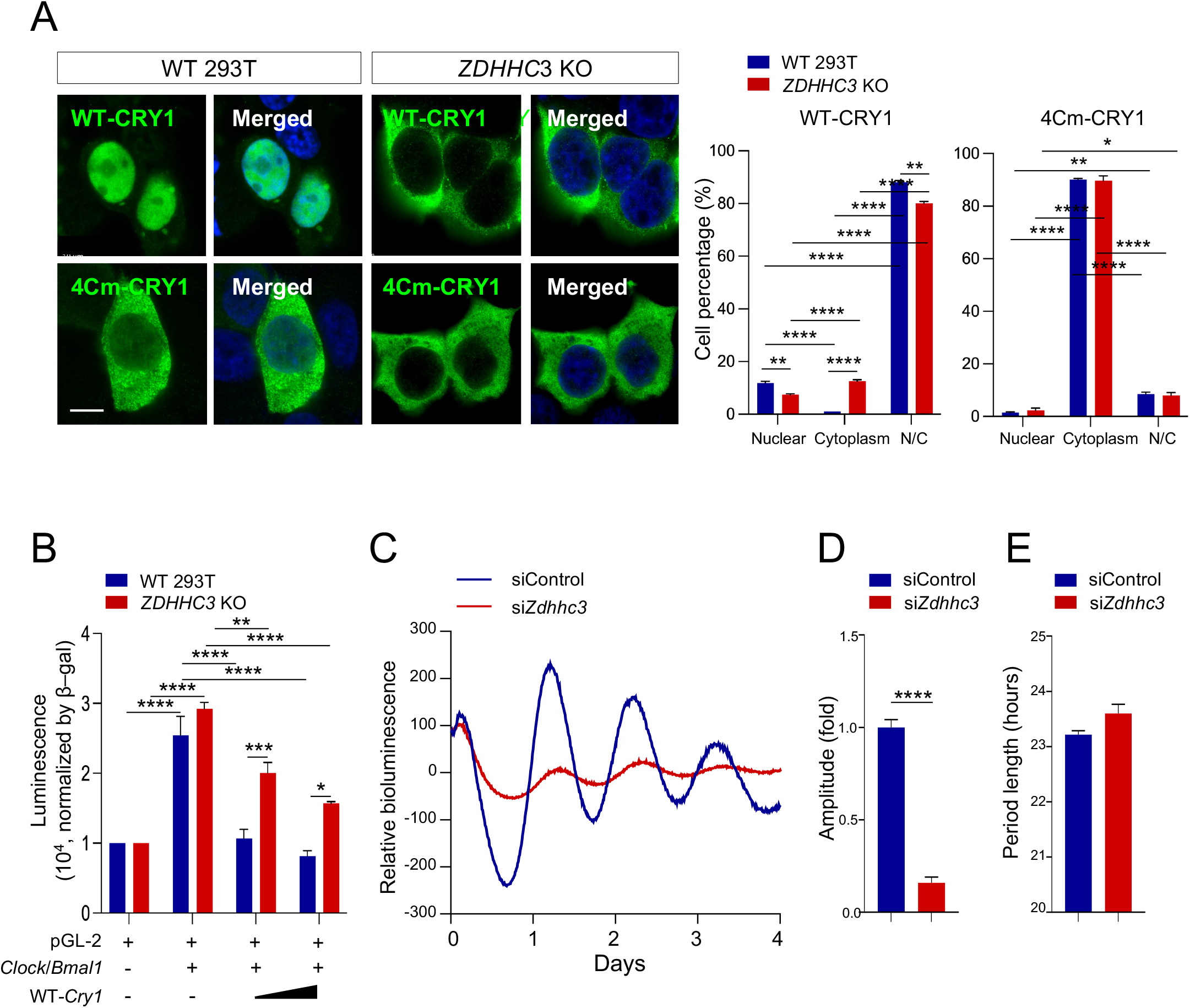
DHHC3 is required for CRY1 localization and cellular circadian rhythms. (**A**) STED microscopy analysis of WT-CRY1 and 4Cm-CRY1 localization in WT and *ZDHHC3* KO 293T cells. Cells were transfected with Flag-tagged WT-Cry1 and 4Cm-Cry1 and imaged using high-resolution STED microscopy with FLAG antibody staining. Right panel: Quantification of the percentage of cells showing nuclear, cytoplasmic, or mixed nuclear/cytoplasmic localization. Data are presented as mean ± SEM (n = 3). \**p* < 0.05, \*\**p* < 0.01 and \*\*\*\**p* < 0.0001; two-way ANOVA indicates statistical differences between groups. Scale bar: 10 μm. (**B**) Comparison of pGL-2 luciferase reporter activity reflecting CLOCK-BMAL1 function in WT and *ZDHHC3* KO 293T cells. 293T cells were transfected with the indicated plasmids, and luciferase activity was measured. Data are presented as mean ± SEM (n = 3–4). \**p* < 0.05, \*\**p* < 0.01, \*\*\**p* < 0.001 and \*\*\*\**p* < 0.0001; Two-way ANOVA indicates statistical differences between groups. (C-E) DHHC3 knockdown reduced circadian amplitude in mouse Per2::LucSV reporter fibroblast cells. Representative PER2::LUC bioluminescence traces of siControl and si*Zdhhc3*-treated cells. (**C**) Average circadian amplitudes (**D**) and period lengths (**E**) in Per2::LucSV fibroblast cells (n = 4/group). \*\*\*\**p* < 0.0001; Student’s *t*-test indicates a significant difference between siControl and si*Zdhhc3*-treated cells.

To examine the functional impact of DHHC3 on CRY1-mediated transcriptional repression, we performed a luciferase reporter assay using WT and *ZDHHC3* KO 293T cells co-transfected with the Per2 E’ box-driven pGL-2 luciferase reporter (Yoo et al., 2005) and WT-CRY1. As expected, CRY1 repressed CLOCK:BMAL1-driven luciferase activity in WT 293T cells (Fig. 5B). However, this repression was significantly attenuated in *ZDHHC3* KO 293T cells (Fig. 5B), indicating that DHHC3-mediated S-acylation is required for optimal CRY1 repressor activity. To evaluate whether DHHC3 influences molecular circadian rhythms, we performed siRNA-mediated knockdown of *Zdhhc3* in Per2::LucSV reporter fibroblast cells and monitored circadian bioluminescence rhythms. *Zdhhc3* knockdown was confirmed by immunoblotting (Fig. S5). Interestingly, depletion of DHHC3 significantly reduced the amplitude of PER2::LUC oscillations, while leaving the period unaffected (Figs. 5C-E). Collectively, these results strongly suggest that DHHC3 plays a critical role in maintaining robust circadian rhythms, likely through its regulation of CRY1 localization and repressor function.

## Discussion

In this study, we identify S-acylation as a novel post-translational modification regulating the function of CRY1, a core component of the mammalian circadian clock. We demonstrate that CRY1 undergoes dynamic S-acylation at multiple cysteine residues, catalyzed primarily by the acyltransferase DHHC3. This lipid modification is essential for CRY1 nuclear localization, stable interaction with clock components such as PER2 and BMAL1, and efficient repression of CLOCK:BMAL1-driven transcription. Loss of CRY1 S-acylation, either through cysteine mutagenesis or DHHC3 deficiency, compromises its repressor function and cellular circadian rhythms. A recent study reported that another core clock protein, CLOCK, is S-acylated by DHHC5 at Cys194 in a human non-small cell lung cancer cell line, leading to enhanced CLOCK protein stability (Peng et al., 2024). In comparison, we did not observe CLOCK S-acylation in our screen of the core clock proteins, potentially due to differences in cell types and detection methods. Regardless, these findings suggest a novel role of reversible protein lipidation in regulating circadian timing in different cell types and establish the DHHC family acyltransferases as a new class of circadian regulatory proteins.

Previous studies have shown that CRY1 is subjected to post-translational modifications including phosphorylation and ubiquitination, which modulate its stability, subcellular localization, and transcriptional activity (Gao et al., 2013; Hirano et al., 2013; Kim et al., 2022; Lamia et al., 2009; Yagita et al., 2002; Yoo et al., 2013). However, the molecular mechanism that governs the rhythmic trafficking of CRY1 between cellular compartments and facilitates its highly dynamic interactions with regulatory partners remains incompletely understood. S-acylation is uniquely suited for this regulatory role, as it induces rapid and reversible changes in protein hydrophobicity, prompting dynamic membrane association and subcellular localization. In other cellular processes, S-acylation has been shown to affect protein-protein interactions at membrane sites such as the plasma membrane and intracellular organelles (Abrami et al., 2008; Linder and Deschenes, 2007; Tewari et al., 2021; Zheng et al., 2023). This concept is further supported by our findings that loss of CRY1 S-acylation impairs not only the formation of the nuclear repressor complex but also interactions with key cytoplasmic binding partners, including PER2 and FBXL21. Moreover, acylation-dependent changes in CRY1 ubiquitination suggest a functional crosstalk between these two modifications. Given that DHHC protein acyltransferases are membrane-bound enzymes, it is plausible that transient membrane association of CRY1 facilitates its S-acylation and enables interactions that control its stability, trafficking, and integration into the core circadian machinery. In contrast to CRY1, we found that CRY2 does not appear to be acylated *in vitro* or *in vivo*, despite the presence of multiple cysteine residues in CRY2. Importantly, the identified cysteine residues required for CRY1 S-acylation (C24, C58, C259, and C315) are not conserved in CRY2, providing a plausible molecular basis for this paralog-specific modification. All four S-acylated cysteines are located within the N-terminal photolyase homology region (PHR) of CRY1 (aa 1–490), which forms the core folded domain of CRY1 mainly responsible for mediating protein–protein interactions (Czarna et al., 2013; Michael et al., 2017; Miller et al., 2021; Todo et al., 1996). The absence of conserved acylation sites in CRY2 is consistent with prior studies demonstrating structural and functional divergence between CRY1 and CRY2 in transcriptional repression and circadian timing (Kriebs et al., 2017; Miller et al., 2021; Rosensweig et al., 2018; van der Horst et al., 1999)

Our biochemical and gain- and loss-of-function studies strongly implicate DHHC3 as the primary acyltransferase responsible for CRY1 S-acylation. Previously, DHHC3 has been linked to the acylation of multiple signaling receptors and regulators, such as GABA receptors (Keller et al., 2004), lamin-binding integrins (Sharma et al., 2012), and G-protein signaling regulator (Wang et al., 2010); our results now extend its role to include regulation of circadian timing. Interestingly, *Zdhhc3* expression exhibits circadian oscillations in multiple tissues, including the suprachiasmatic nucleus (SCN)(Mure et al., 2018; Wen et al., 2020), and its promoter shows strong binding by core clock transcription factors (Zou et al., 2024). These findings suggest that DHHC3, and potentially other members of the DHHC family, may be controlled by the circadian clock, forming a regulatory feedback loop between the S-acylation machinery and the circadian oscillator.

The DHHC enzyme family comprises at least 23 members, many of which are broadly expressed and share overlapping substrate specificity (Fukata et al., 2006; Greaves and Chamberlain, 2011). In addition to DHHC3, our data also revealed interactions between CRY1 and other DHHC enzymes, including DHHC17, DHHC18, DHHC19, and DHHC22 (Fig. S4), suggesting that CRY1 may be S-acylated by multiple acyltransferases. This raises the possibility that distinct DHHC enzymes may modify CRY1 in a tissue-specific manner or under conditions where DHHC3 expression is low or absent. Furthermore, because S-acylation is a reversible modification, S-acylation of CRY1 may also be dynamically modulated by acyl-protein thioesterases (APTs) that remove the fatty acid moiety, potentially in response to circadian or metabolic cues (Lin and Conibear, 2015). However, the APTs responsible for CRY1 de-acylation remain to be identified, which may reveal another novel class of regulatory enzymes in the context of circadian biology. Thus, future studies are necessary to fully elucidate the enzymatic machinery governing reversible S-acylation of clock proteins, including CRY1, and to uncover their broader clock-controlled physiological relevance.

In summary, our findings establish S-acylation as a novel regulatory mechanism within the mammalian circadian clock and identify DHHC3 as a key enzymatic modulator of CRY1 function. This work expands the current landscape of post-translational regulation in circadian biology and highlights protein S-acylation as a critical signaling node that links CRY1 intracellular localization, protein-protein interactions, and transcriptional repression activity. From a translational perspective, the reversibility and enzymatic specificity of S-acylation render this modification an attractive target for pharmacological modulation of circadian timing in disorders associated with clock misalignment, such as shift work–related sleep disturbances, metabolic syndrome, and other age-associated chronic diseases.

## Material & Methods

### Animal Studies and Ethics Statement

Eight-week-old male C57BL/6J mice (Stock #000664; Jackson Laboratory, Bar Harbor, ME, USA) were obtained and housed under a 12 h light/12 h dark cycle (LD 12:12). All experimental procedures conducted in accordance with protocols approved by the Center for Laboratory Animal Medicine and Care (CLAMC) at The University of Texas Health Science Center at Houston (UTHealth Houston).

### Cell Culture and Transfection

HEK293T (ATCC CRL-3216) cells were maintained in Dulbecco’s Modified Eagle’s Medium (DMEM) supplemented with 10% fetal bovine serum (FBS; GenDEPOT, Houston, TX, USA). For immunoprecipitation experiments, 2 × 10^6^ cells were plated in 60-mm dishes, and plasmids were delivered using iMFectin (GenDEPOT) following the manufacturer’s protocol.

### siRNA Transfection and real-time bioluminescence measurements

Mouse *Per2::LucSV* reporter fibroblast cells were plated in 35 mm plates in DMEM supplemented with 10% FBS and 1% penicillin/streptomycin, as previously described (Yoo et al., 2017; Yoo et al., 2004). For *Zdhhc3* knockdown, cells were transfected with a scrambled control siRNA or 50 nM of MISSION siRNAs targeting murine *Zdhhc3* (ZDHHC3; SASI_Mm02_00331193, Sigma-Aldrich, St. Louis, MO, USA) using Lipofectamine RNAiMAX (ThermoFisher Scientific) following the manufacturer’s instructions. Thirty-six hours post-transfection, the cells were treated with 200 nM dexamethasone (Dex) for 1 hour, then cultured with luciferin-containing recording media and monitored continuously for bioluminescence over 6 days using a LumiCycle luminometer (Actimetrics) for continuous bioluminescence monitoring. Bioluminescence data were analyzed by the LumiCycle data analysis program (Actimetrics).

### Acyl-RAC assay

The Acyl-RAC assay was performed as previously described (Kodakandla et al., 2022). Briefly, cell and tissue lysates were prepared in PBS containing 1% dodecyl β-D-maltoside, supplemented with a protease and phosphatase inhibitor cocktail (GenDEPOT), the acyl protein thioesterase inhibitor ML211 (Cayman Chemical), and the serine protease inhibitor PMSF (10 mM). Protein concentrations were assessed using a BCA assay. 500 ug of protein lysates were precipitated using a 2:1 methanol–chloroform mixture. Following centrifugation, the resulting protein pellets were incubated with 0.2% methyl methanethiosulfonate for 15 min at 42 °C. To remove residual methyl methanethiosulfonate, methanol–chloroform precipitation was repeated three times. The final protein pellets were resuspended in 2SHB buffer (2% SDS, 5 mM EDTA, 100 mM HEPES, pH 7.4) and incubated with 400 mM hydroxylamine (HA) to cleave thioester bonds. Thiol-Sepharose resin was then added, and the samples were incubated overnight at 4 °C. Bound proteins were eluted with 10 mM DTT in SDS sample buffer (1% SDS, 50 mM Tris–HCl, 10% glycerol, and 1% bromophenol blue) and subsequently analyzed by immunoblotting.

### Generation of *ZDHHC3* CRISPR KO cell lines

*ZDHHC3* KO 293T cells were generated as previously described (Lim et al., 2022; Wirianto et al., 2020). Briefly, pairs of guide RNAs targeting human *ZDHHC3* were designed and cloned into the BsmBI site of the LentiCRISPR v2 vector. The guide RNA sequences were as follows: *ZDHHC3*(-) forward, CACCGTGTGGTTTATCCGTGA; reverse, AAACTCACGGATAAACCACA; human *ZDHHC3*(+) forward, CACCGACTCTCGATGAATTCTTTAG; reverse, AAACCTAAAGAATTCATCGAGAGT. HEK293T cells were transfected with the resulting constructs and selected with puromycin (2 μg/ml). Successful *ZDHHC3* knockout was confirmed by qPCR.

### Reporter assays

Reporter assays were carried out as previously described (Yoo et al., 2017). Briefly, WT or *ZDHHC3* KO 293T cells were seeded in 12-well plates at 2 x 10^5^ cells per well. The following day, cells were transfected with the indicated plasmids using IMFectin (GenDEPOT). After 36 hours, the cells were lysed, and luminescence was measured using a Tecan Spark 10M microplate reader (TECAN, Männedorf, Switzerland).

### Identification of CRY1 acylation sites by LC-MS/MS and generation of the 4Cm-CRY1 mutant

To increase protein abundance, we transfected WT 293T cells with Strep-tagged CRY1 (IBA Strep-tag system). Following cell lysis and hydroxylamine treatment, S-acylated proteins were reacted with thiol-reactive resin. Unlike the standard Acyl-RAC workflow, immobilized proteins were digested directly on the beads using trypsin, and the unbound peptides were removed by extensive washing. The retained peptides were then eluted by reducing disulfide bonds using DTT and subjected to LC-MS/MS analysis. To investigate the S-acylation sites in CRY1, peptide sequences from DTT elution fraction were analyzed, and we identified Cys24, Cys58, Cys259, and Cys315, as potential S-acylation sites in CRY1. A 4Cm-Cry1 mutant plasmid, in which Cys24, Cys58, Cys259, and Cys315 were mutated to alanine, was synthesized by GenScript (Piscataway, NJ, USA). Structural and computational analyses of 4Cm-CRY1 were conducted using AlphaFold3 structural modeling (Abramson et al., 2024) and Rosetta-based mutational energy analysis (Alford et al., 2017; Kellogg et al., 2011).

### Immunoblotting, Immunoprecipitation, and Immunofluorescence staining

Immunoblotting, immunoprecipitation, and immunofluorescence staining were carried out as previously described (Wirianto et al., 2020; Yoo et al., 2013). To assess protein degradation rates and ubiquitination, HEK293T cells were transfected with expression constructs encoding pCMV10-3XFlag-WT-Cry1, pCMV10-3XFlag-4Cm-Cry1, HA-Fbxl3, HA-Fbxl21, as well as untagged Fbxl3 and Fbxl21, and HA-Ub. For cycloheximide (CHX) chase assays, after thirty-six hours, CHX (100 μg/mL) was added, and cells were collected at sequential time points for analysis. To calculate protein half-lives, nonlinear one-phase decay fitting was used in GraphPad Prism. Ubiquitination assays were conducted as previously described (Wirianto et al., 2020; Yoo et al., 2013). Antibodies against Flag-HRP (Sigma-Aldrich), anti-HA (Roche), and anti-V5 (Thermo Fisher Scientific) were used to detect ectopically expressed proteins by immunoblotting. Antibodies against DHHC3 (ThermoFisher Scientific), Calnexin (ProteinTech), and GAPDH (Abclonal Biotechnology) were used to detect endogenous proteins. Antibodies against CRY1 (epitope: aa 496-606) and CRY2 (epitope: aa 514-592) were generated in guinea pigs (Cocalico Biologicals) and purified by serum affinity chromatography (Yoo et al., 2013). For STED imaging, anti-FLAG antibody (Sigma) and goat anti-mouse STAR ORANGE secondary antibody were used. Images were acquired using a STELLARIS confocal microscope (Leica Microsystems) or STED microscope (Abberior Instruments, Göttingen, Germany).

## Acknowledgments

We thank the Center for Advanced Microscopy of the UTHealth McGovern Medical School for technical assistance.

## Author contributions

S.Y. and A.A. designed the study. J.L., S.B., C.H., S.Y.K., M.L.B. and S.Y.J. performed experiments and analyzed the results. S.Y., A.A., and J. L. drafted the manuscript. T.M., K.L.T, H.K.L. S.Y.J. and Z.C. reviewed and revised the manuscript.

## References

Abrami, L., B. Kunz, I. Iacovache, and F.G. van der Goot. 2008. Palmitoylation and ubiquitination regulate exit of the Wnt signaling protein LRP6 from the endoplasmic reticulum. Proc Natl Acad Sci U S A. 105:5384–5389.

Abramson, J., J. Adler, J. Dunger, R. Evans, T. Green, A. Pritzel, O. Ronneberger, L. Willmore, A.J. Ballard, J. Bambrick, S.W. Bodenstein, D.A. Evans, C.C. Hung, M. O’Neill, D. Reiman, K. Tunyasuvunakool, Z. Wu, A. Zemgulyte, E. Arvaniti, C. Beattie, O. Bertolli, A. Bridgland, A. Cherepanov, M. Congreve, A.I. Cowen-Rivers, A. Cowie, M. Figurnov, F.B. Fuchs, H. Gladman, R. Jain, Y.A. Khan, C.M.R. Low, K. Perlin, A. Potapenko, P. Savy, S. Singh, A. Stecula, A. Thillaisundaram, C. Tong, S. Yakneen, E.D. Zhong, M. Zielinski, A. Zidek, V. Bapst, P. Kohli, M. Jaderberg, D. Hassabis, and J.M. Jumper. 2024. Accurate structure prediction of biomolecular interactions with AlphaFold 3. Nature. 630:493–500.

Alford, R.F., A. Leaver-Fay, J.R. Jeliazkov, M.J. O’Meara, F.P. DiMaio, H. Park, M.V. Shapovalov, P.D. Renfrew, V.K. Mulligan, K. Kappel, J.W. Labonte, M.S. Pacella, R. Bonneau, P. Bradley, R.L. Dunbrack, Jr., R. Das, D. Baker, B. Kuhlman, T. Kortemme, and J.J. Gray. 2017. The Rosetta All-Atom Energy Function for Macromolecular Modeling and Design. J Chem Theory Comput. 13:3031–3048.

Anwar, M.U., and F.G. van der Goot. 2023. Refining S-acylation: Structure, regulation, dynamics, and therapeutic implications. J Cell Biol. 222.

Bass, J., and M.A. Lazar. 2016. Circadian time signatures of fitness and disease. Science. 354:994–999.

Bass, J., and J.S. Takahashi. 2010. Circadian integration of metabolism and energetics. Science. 330:1349–1354.

Cederroth, C.R., U. Albrecht, J. Bass, S.A. Brown, J. Dyhrfjeld-Johnsen, F. Gachon, C.B. Green, M.H. Hastings, C. Helfrich-Forster, J.B. Hogenesch, F. Levi, A. Loudon, G.B. Lundkvist, J.H. Meijer, M. Rosbash, J.S. Takahashi, M. Young, and B. Canlon. 2019. Medicine in the Fourth Dimension. Cell Metab. 30:238–250.

Chamberlain, L.H., and M.J. Shipston. 2015. The physiology of protein S-acylation. Physiological reviews. 95:341–376.

Chaves, I., K. Yagita, S. Barnhoorn, H. Okamura, G.T. van der Horst, and F. Tamanini. 2006. Functional evolution of the photolyase/cryptochrome protein family: importance of the C terminus of mammalian CRY1 for circadian core oscillator performance. Mol Cell Biol. 26:1743–1753.

Chen, B., Y. Sun, J. Niu, G.K. Jarugumilli, and X. Wu. 2018. Protein Lipidation in Cell Signaling and Diseases: Function, Regulation, and Therapeutic Opportunities. Cell Chem Biol. 25:817–831.

Chen, J.J., Y. Fan, and D. Boehning. 2021. Regulation of Dynamic Protein S-Acylation. Front Mol Biosci. 8:656440.

Czarna, A., A. Berndt, H.R. Singh, A. Grudziecki, A.G. Ladurner, G. Timinszky, A. Kramer, and E. Wolf. 2013. Structures of Drosophila cryptochrome and mouse cryptochrome1 provide insight into circadian function. Cell. 153:1394–1405.

F, S.M., L. Abrami, M.E. Linder, S.X. Bamji, B.C. Dickinson, and F.G. van der Goot. 2024. Mechanisms and functions of protein S-acylation. Nat Rev Mol Cell Biol. 25:488–509.

Fukata, Y., T. Iwanaga, and M. Fukata. 2006. Systematic screening for palmitoyl transferase activity of the DHHC protein family in mammalian cells. Methods. 40:177–182.

Gallego, M., and D.M. Virshup. 2007. Post-translational modifications regulate the ticking of the circadian clock. Nat Rev Mol Cell Biol. 8:139–148.

Gao, P., S.H. Yoo, K.J. Lee, C. Rosensweig, J.S. Takahashi, B.P. Chen, and C.B. Green. 2013. Phosphorylation of the cryptochrome 1 C-terminal tail regulates circadian period length. The Journal of biological chemistry. 288:35277–35286.

Greaves, J., and L.H. Chamberlain. 2011. DHHC palmitoyl transferases: substrate interactions and (patho)physiology. Trends Biochem Sci. 36:245–253.

Green, C.B., J.S. Takahashi, and J. Bass. 2008. The meter of metabolism. Cell. 134:728–742.

Hirano, A., Y.H. Fu, and L.J. Ptacek. 2016. The intricate dance of post-translational modifications in the rhythm of life. Nature structural & molecular biology. 23:1053–1060.

Hirano, A., K. Yumimoto, R. Tsunematsu, M. Matsumoto, M. Oyama, H. Kozuka-Hata, T. Nakagawa, D. Lanjakornsiripan, K.I. Nakayama, and Y. Fukada. 2013. FBXL21 regulates oscillation of the circadian clock through ubiquitination and stabilization of cryptochromes. Cell. 152:1106–1118.

Hughes, M.E., J.B. Hogenesch, and K. Kornacker. 2010. JTK_CYCLE: an efficient nonparametric algorithm for detecting rhythmic components in genome-scale data sets. J Biol Rhythms. 25:372–380.

Jiang, H., X. Zhang, X. Chen, P. Aramsangtienchai, Z. Tong, and H. Lin. 2018. Protein Lipidation: Occurrence, Mechanisms, Biological Functions, and Enabling Technologies. Chem Rev. 118:919–988.

Keller, C.A., X. Yuan, P. Panzanelli, M.L. Martin, M. Alldred, M. Sassoe-Pognetto, and B. Luscher. 2004. The gamma2 subunit of GABA(A) receptors is a substrate for palmitoylation by GODZ. J Neurosci. 24:5881–5891.

Kellogg, E.H., A. Leaver-Fay, and D. Baker. 2011. Role of conformational sampling in computing mutation-induced changes in protein structure and stability. Proteins. 79:830–838.

Kim, Y.Y., H. Jang, G. Lee, Y.G. Jeon, J.H. Sohn, J.S. Han, W.T. Lee, J. Park, J.Y. Huh, H. Nahmgoong, S.M. Han, J. Kim, M. Pak, S. Kim, J.S. Kim, and J.B. Kim. 2022. Hepatic GSK3beta-Dependent CRY1 Degradation Contributes to Diabetic Hyperglycemia. Diabetes. 71:1373–1387.

Kodakandla, G., S.J. West, Q. Wang, R. Tewari, M.X. Zhu, A.M. Akimzhanov, and D. Boehning. 2022. Dynamic S-acylation of the ER-resident protein stromal interaction molecule 1 (STIM1) is required for store-operated Ca(2+) entry. The Journal of biological chemistry. 298:102303.

Kriebs, A., S.D. Jordan, E. Soto, E. Henriksson, C.R. Sandate, M.E. Vaughan, A.B. Chan, D. Duglan, S.J. Papp, A.L. Huber, M.E. Afetian, R.T. Yu, X. Zhao, M. Downes, R.M. Evans, and K.A. Lamia. 2017. Circadian repressors CRY1 and CRY2 broadly interact with nuclear receptors and modulate transcriptional activity. Proc Natl Acad Sci U S A. 114:8776–8781.

Kume, K., M.J. Zylka, S. Sriram, L.P. Shearman, D.R. Weaver, X. Jin, E.S. Maywood, M.H. Hastings, and S.M. Reppert. 1999. mCRY1 and mCRY2 are essential components of the negative limb of the circadian clock feedback loop. Cell. 98:193–205.

Kurabayashi, N., T. Hirota, M. Sakai, K. Sanada, and Y. Fukada. 2010. DYRK1A and glycogen synthase kinase 3beta, a dual-kinase mechanism directing proteasomal degradation of CRY2 for circadian timekeeping. Mol Cell Biol. 30:1757–1768.

Lamia, K.A., U.M. Sachdeva, L. DiTacchio, E.C. Williams, J.G. Alvarez, D.F. Egan, D.S. Vasquez, H. Juguilon, S. Panda, R.J. Shaw, C.B. Thompson, and R.M. Evans. 2009. AMPK regulates the circadian clock by cryptochrome phosphorylation and degradation. Science. 326:437–440.

Lim, J.Y., E. Kim, C.M. Douglas, M. Wirianto, C. Han, K. Ono, S.Y. Kim, J.H. Ji, C.K. Tran, Z. Chen, K.A. Esser, and S.H. Yoo. 2022. The circadian E3 ligase FBXL21 regulates myoblast differentiation and sarcomere architecture via MYOZ1 ubiquitination and NFAT signaling. PLoS Genet. 18:e1010574.

Lin, D.T., and E. Conibear. 2015. ABHD17 proteins are novel protein depalmitoylases that regulate N-Ras palmitate turnover and subcellular localization. Elife. 4:e11306.

Linder, M.E., and R.J. Deschenes. 2007. Palmitoylation: policing protein stability and traffic. Nat Rev Mol Cell Biol. 8:74–84.

Masri, S., and P. Sassone-Corsi. 2018. The emerging link between cancer, metabolism, and circadian rhythms. Nat Med. 24:1795–1803.

Michael, A.K., J.L. Fribourgh, Y. Chelliah, C.R. Sandate, G.L. Hura, D. Schneidman-Duhovny, S.M. Tripathi, J.S. Takahashi, and C.L. Partch. 2017. Formation of a repressive complex in the mammalian circadian clock is mediated by the secondary pocket of CRY1. Proc Natl Acad Sci U S A. 114:1560–1565.

Miller, S., A. Srivastava, Y. Nagai, Y. Aikawa, F. Tama, and T. Hirota. 2021. Structural differences in the FAD-binding pockets and lid loops of mammalian CRY1 and CRY2 for isoform-selective regulation. Proc Natl Acad Sci U S A. 118.

Millius, A., K.L. Ode, and H.R. Ueda. 2019. A period without PER: understanding 24-hour rhythms without classic transcription and translation feedback loops. F1000Res. 8.

Mitchell, D.A., A. Vasudevan, M.E. Linder, and R.J. Deschenes. 2006. Protein palmitoylation by a family of DHHC protein S-acyltransferases. Journal of lipid research. 47:1118–1127.

Miyazaki, K., M. Mesaki, and N. Ishida. 2001. Nuclear entry mechanism of rat PER2 (rPER2): role of rPER2 in nuclear localization of CRY protein. Mol Cell Biol. 21:6651–6659.

Mure, L.S., H.D. Le, G. Benegiamo, M.W. Chang, L. Rios, N. Jillani, M. Ngotho, T. Kariuki, O. Dkhissi-Benyahya, H.M. Cooper, and S. Panda. 2018. Diurnal transcriptome atlas of a primate across major neural and peripheral tissues. Science. 359.

Okamoto-Uchida, Y., J. Izawa, A. Nishimura, A. Hattori, N. Suzuki, and J. Hirayama. 2019. Posttranslational Modifications are Required for Circadian Clock Regulation in Vertebrates. Curr Genomics. 20:332–339.

Parico, G.C.G., I. Perez, J.L. Fribourgh, B.N. Hernandez, H.W. Lee, and C.L. Partch. 2020. The human CRY1 tail controls circadian timing by regulating its association with CLOCK:BMAL1. Proc Natl Acad Sci U S A. 117:27971–27979.

Peng, F., J. Lu, K. Su, X. Liu, H. Luo, B. He, C. Wang, X. Zhang, F. An, D. Lv, Y. Luo, Q. Su, T. Jiang, Z. Deng, B. He, L. Xu, T. Guo, J. Xiang, C. Gu, L. Wang, G. Xu, Y. Xu, M. Li, K.W. Kelley, B. Cui, and Q. Liu. 2024. Oncogenic fatty acid oxidation senses circadian disruption in sleep-deficiency-enhanced tumorigenesis. Cell Metab. 36:1598–1618 e1511.

Rosensweig, C., K.A. Reynolds, P. Gao, I. Laothamatas, Y. Shan, R. Ranganathan, J.S. Takahashi, and C.B. Green. 2018. An evolutionary hotspot defines functional differences between CRYPTOCHROMES. Nature communications. 9:1138.

Sharma, C., I. Rabinovitz, and M.E. Hemler. 2012. Palmitoylation by DHHC3 is critical for the function, expression, and stability of integrin alpha6beta4. Cell Mol Life Sci. 69:2233–2244.

Siepka, S.M., S.H. Yoo, J. Park, W. Song, V. Kumar, Y. Hu, C. Lee, and J.S. Takahashi. 2007. Circadian mutant Overtime reveals F-box protein FBXL3 regulation of cryptochrome and period gene expression. Cell. 129:1011–1023.

Takahashi, J.S. 2017. Transcriptional architecture of the mammalian circadian clock. Nat Rev Genet. 18:164–179.

Tamanini, F., K. Yagita, H. Okamura, and G.T. van der Horst. 2005. Nucleocytoplasmic shuttling of clock proteins. Methods in enzymology. 393:418–435.

Tewari, R., B. Shayahati, Y. Fan, and A.M. Akimzhanov. 2021. T cell receptor-dependent S-acylation of ZAP-70 controls activation of T cells. The Journal of biological chemistry. 296:100311.

Tewari, R., S.J. West, B. Shayahati, and A.M. Akimzhanov. 2020. Detection of Protein S-Acylation using Acyl-Resin Assisted Capture. Journal of visualized experiments: JoVE.

Todo, T., H. Ryo, K. Yamamoto, H. Toh, T. Inui, H. Ayaki, T. Nomura, and M. Ikenaga. 1996. Similarity among the Drosophila (6-4)photolyase, a human photolyase homolog, and the DNA photolyase-blue-light photoreceptor family. Science. 272:109–112.

Tsutsumi, R., Y. Fukata, J. Noritake, T. Iwanaga, F. Perez, and M. Fukata. 2009. Identification of G protein alpha subunit-palmitoylating enzyme. Mol Cell Biol. 29:435–447.

van der Horst, G.T., M. Muijtjens, K. Kobayashi, R. Takano, S. Kanno, M. Takao, J. de Wit, A. Verkerk, A.P. Eker, D. van Leenen, R. Buijs, D. Bootsma, J.H. Hoeijmakers, and A. Yasui. 1999. Mammalian Cry1 and Cry2 are essential for maintenance of circadian rhythms. Nature. 398:627–630.

van der Schalie, E.A., F.E. Conte, K.E. Marz, and C.B. Green. 2007. Structure/function analysis of Xenopus cryptochromes 1 and 2 reveals differential nuclear localization mechanisms and functional domains important for interaction with and repression of CLOCK-BMAL1. Mol Cell Biol. 27:2120–2129.

Vitaterna, M.H., C.P. Selby, T. Todo, H. Niwa, C. Thompson, E.M. Fruechte, K. Hitomi, R.J. Thresher, T. Ishikawa, J. Miyazaki, J.S. Takahashi, and A. Sancar. 1999. Differential regulation of mammalian period genes and circadian rhythmicity by cryptochromes 1 and 2. Proc Natl Acad Sci U S A. 96:12114–12119.

Wang, J., Y. Xie, D.W. Wolff, P.W. Abel, and Y. Tu. 2010. DHHC protein-dependent palmitoylation protects regulator of G-protein signaling 4 from proteasome degradation. FEBS Lett. 584:4570–4574.

Wen, S., D. Ma, M. Zhao, L. Xie, Q. Wu, L. Gou, C. Zhu, Y. Fan, H. Wang, and J. Yan. 2020. Spatiotemporal single-cell analysis of gene expression in the mouse suprachiasmatic nucleus. Nature neuroscience. 23:456–467.

West, S.J., D. Boehning, and A.M. Akimzhanov. 2022. Regulation of T cell function by protein S-acylation. Front Physiol. 13:1040968.

Wirianto, M., J. Yang, E. Kim, S. Gao, K.R. Paudel, J.M. Choi, J. Choe, G.F. Gloston, P. Ademoji, R. Parakramaweera, J. Jin, K.A. Esser, S.Y. Jung, Y.J. Geng, H.K. Lee, Z. Chen, and S.H. Yoo. 2020. The GSK-3beta-FBXL21 Axis Contributes to Circadian TCAP Degradation and Skeletal Muscle Function. Cell reports. 32:108140.

Yagita, K., F. Tamanini, M. Yasuda, J.H. Hoeijmakers, G.T. van der Horst, and H. Okamura. 2002. Nucleocytoplasmic shuttling and mCRY-dependent inhibition of ubiquitylation of the mPER2 clock protein. Embo J. 21:1301–1314.

Yoo, S.H., C.H. Ko, P.L. Lowrey, E.D. Buhr, E.J. Song, S. Chang, O.J. Yoo, S. Yamazaki, C. Lee, and J.S. Takahashi. 2005. A noncanonical E-box enhancer drives mouse Period2 circadian oscillations in vivo. Proc Natl Acad Sci U S A. 102:2608–2613.

Yoo, S.H., S. Kojima, K. Shimomura, N. Koike, E.D. Buhr, T. Furukawa, C.H. Ko, G. Gloston, C. Ayoub, K. Nohara, B.A. Reyes, Y. Tsuchiya, O.J. Yoo, K. Yagita, C. Lee, Z. Chen, S. Yamazaki, C.B. Green, and J.S. Takahashi. 2017. Period2 3’-UTR and microRNA-24 regulate circadian rhythms by repressing PERIOD2 protein accumulation. Proc Natl Acad Sci U S A. 114:E8855–E8864.

Yoo, S.H., J.A. Mohawk, S.M. Siepka, Y. Shan, S.K. Huh, H.K. Hong, I. Kornblum, V. Kumar, N. Koike, M. Xu, J. Nussbaum, X. Liu, Z. Chen, Z.J. Chen, C.B. Green, and J.S. Takahashi. 2013. Competing E3 Ubiquitin Ligases Govern Circadian Periodicity by Degradation of CRY in Nucleus and Cytoplasm. Cell. 152:1091–1105.

Yoo, S.H., S. Yamazaki, P.L. Lowrey, K. Shimomura, C.H. Ko, E.D. Buhr, S.M. Siepka, H.K. Hong, W.J. Oh, O.J. Yoo, M. Menaker, and J.S. Takahashi. 2004. PERIOD2::LUCIFERASE real-time reporting of circadian dynamics reveals persistent circadian oscillations in mouse peripheral tissues. Proc Natl Acad Sci U S A. 101:5339–5346.

Zaballa, M.E., and F.G. van der Goot. 2018. The molecular era of protein S-acylation: spotlight on structure, mechanisms, and dynamics. Crit Rev Biochem Mol Biol. 53:420–451.

Zhang, R., N.F. Lahens, H.I. Ballance, M.E. Hughes, and J.B. Hogenesch. 2014. A circadian gene expression atlas in mammals: implications for biology and medicine. Proceedings of the National Academy of Sciences of the United States of America. 111:16219–16224.

Zheng, S., X. Que, S. Wang, Q. Zhou, X. Xing, L. Chen, C. Hou, J. Ma, P. An, Y. Peng, Y. Yao, Q. Song, J. Li, P. Zhang, and H. Pei. 2023. ZDHHC5-mediated NLRP3 palmitoylation promotes NLRP3-NEK7 interaction and inflammasome activation. Mol Cell. 83:4570–4585 e4577.

Zou, Z., T. Ohta, and S. Oki. 2024. ChIP-Atlas 3.0: a data-mining suite to explore chromosome architecture together with large-scale regulome data. Nucleic Acids Res. 52:W45–W53.

