## supplemental materials for "DHHC3-dependent S-Acylation of CRY1 regulates its subcellular localization and repressor function in the circadian clock"

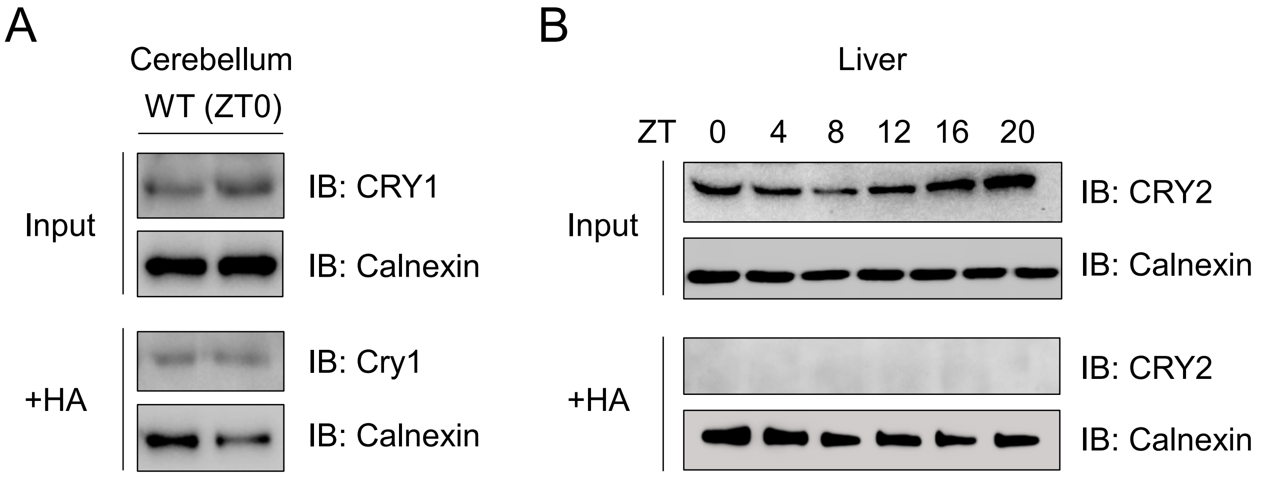


**Fig. S1. S-acylation of CRY1 in cerebellum and S-acylation of CRY2 in the liver.** (**A**) Immunoblot analysis of CRY1-S-acylated protein in the cerebellum of WT mice collected at ZT (Zeitgeber time) 0. Calnexin was used as a loading control. (**B**) Immunoblot analysis of CRY2 S-acylation in the liver tissues of WT C57BL/6J mice collected at the indicated ZT. Calnexin was used as a loading control.

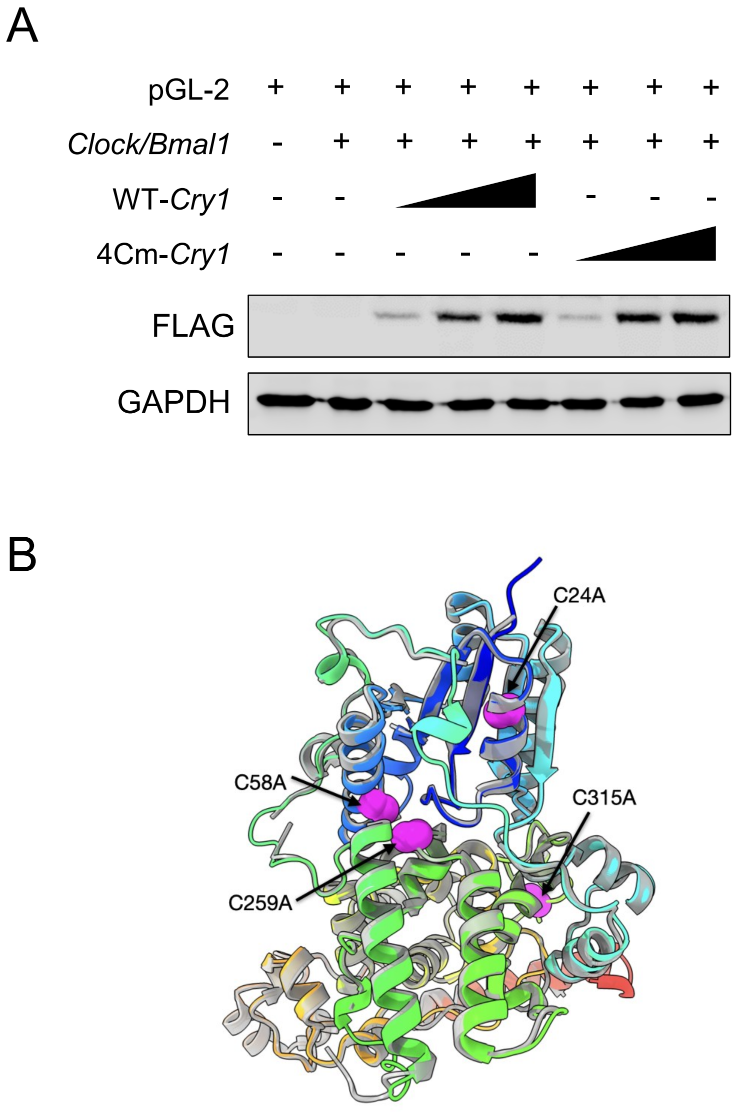


**Fig. S2. CRY1 protein levels for Fig. 2E and structural comparison of WT-CRY1 and 4Cm-CRY1.** (**A**) Immunoblot analysis of CRY1 for Fig. 2E. 293T cells were transfected with constructs as indicated. GAPDH was used as a loading control. (**B**) AlphaFold3 model colored from N- (blue) to C-terminus (red); mutated residues (C24, C58, C259, C315) shown in magenta (spheres). WT CRY1 structure (PDB: 4K0R) is rendered in transparent gray. Structural alignment yields an RMSD of 0.5 Å, indicating a high degree of structural similarity and preservation of the overall fold.

*Fig. S3*

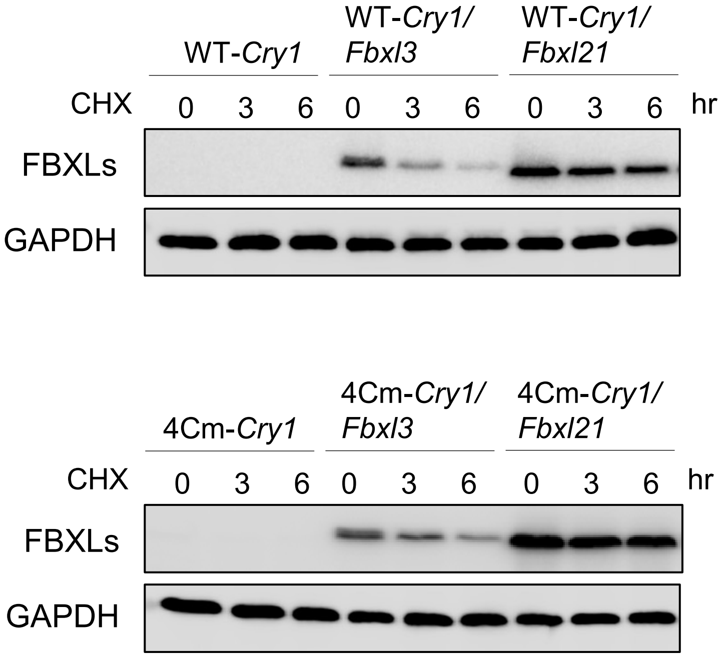


**Fig. S3. FBXL3 and FBXL21 protein levels for Fig. 3B.** Immunoblot analysis of FBXL3 and FBXL21 for Fig. 3B. 293T cells were transfected with flag-tagged WT-*Cry1* and 4Cm-*Cry1* and HA-tagged *Fbxl3* and *Fbxl21* constructs as indicated. Immunoblotting assay was performed using HA antibody. GAPDH was used as a loading control.

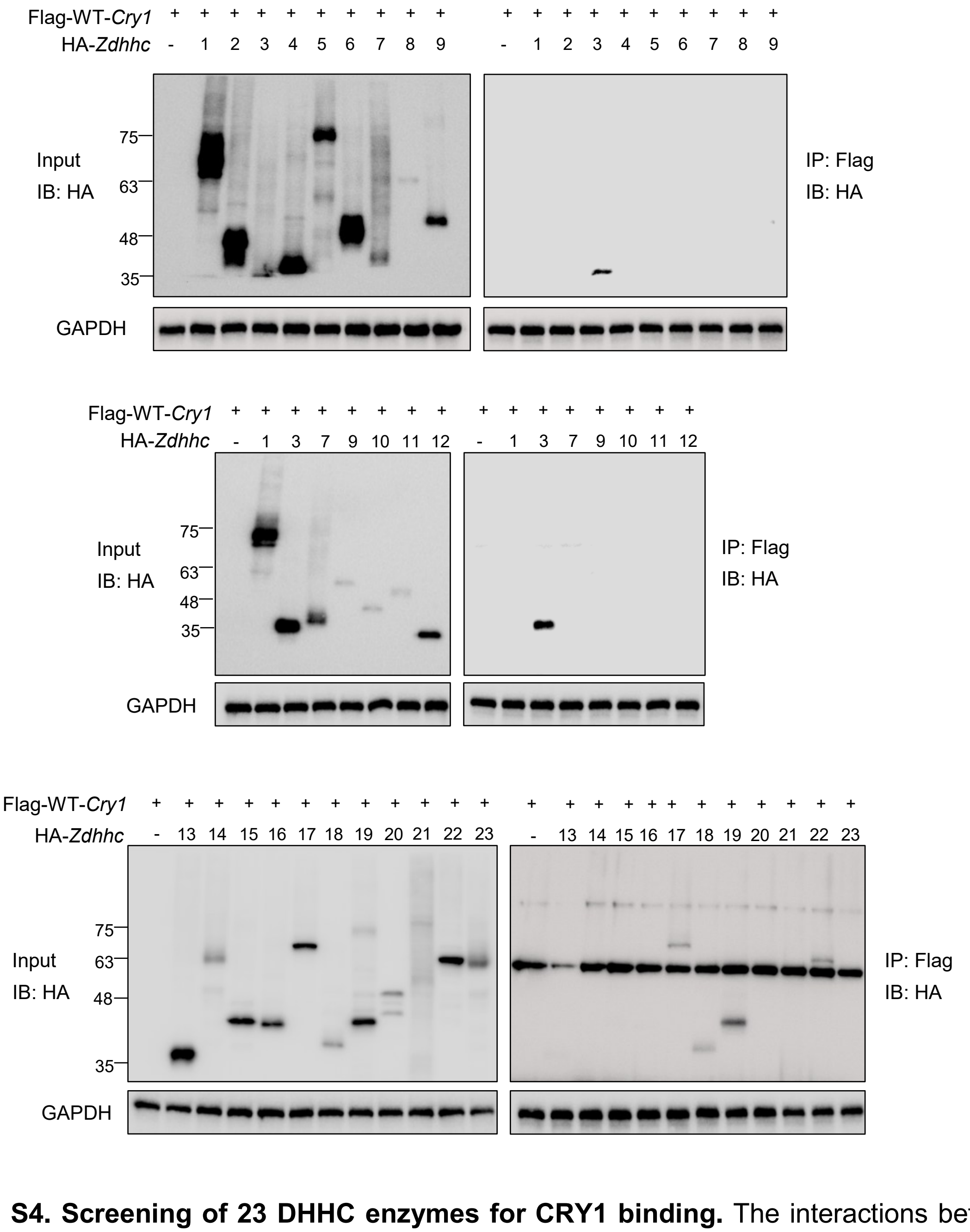


**Fig. S4. Screening of 23 DHHC enzymes for CRY1 binding.** The interactions between 23 DHHC enzymes and WT-CRY1 were analyzed. 293T cells were transfected with the indicated plasmids, and protein complexes were pulled down using anti-Flag beads. GAPDH was used as a loading control.

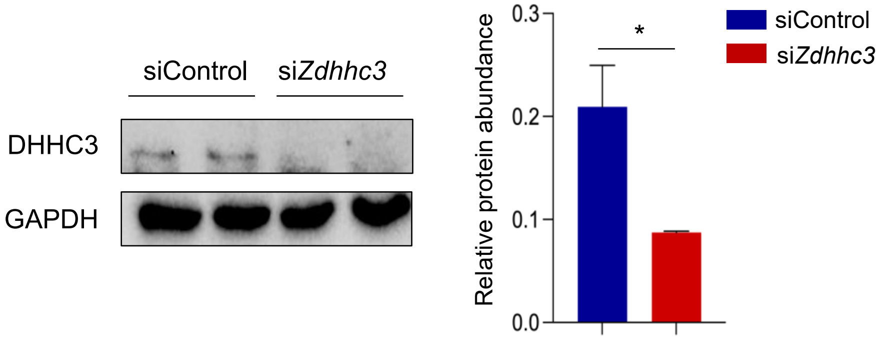


**Fig. S5. DHHC3 knockdown in Per2::LucSV reporter fibroblast cells for Fig. 5C-E.** Immunoblotting analysis of DHHC3 in Per2::LucSV reporter fibroblast cells treated with siControl and si*Zdhhc3*. Data are presented as ± SEM (n = 3). *$p < 0.05$; Student's *t*-test indicates a significant difference between siControl and si*Zdhhc3*-treated cells.
